# Alternative substitutions of N332 in HIV-1_AD8_ gp120 differentially affect envelope glycoprotein function and viral sensitivity to broadly neutralizing antibodies targeting the V3-glycan

**DOI:** 10.1101/2023.11.20.567910

**Authors:** Jeffy Jeffy, Durgadevi Parthasarathy, Shamim Ahmed, Héctor Cervera-Benet, Ulahn Xiong, Miranda Harris, Dmitriy Mazurov, Stephanie Pickthorn, Alon Herschhorn

**Affiliations:** Division of Infectious Diseases and International Medicine, Department of Medicine, University of Minnesota, Minneapolis, Minnesota 55455, USA; Institute for Molecular Virology, University of Minnesota, Minneapolis, MN 55455, USA; Institute of Engineering and Medicine, University of Minnesota, Minneapolis, MN 55455, USA; Center of Genomic Engineering, University of Minnesota, Minneapolis, MN 55455, USA; Microbiology, Immunology, and Cancer Biology Graduate Program, the College of Veterinary Medicine Graduate Program, and Molecular Pharmacology and Therapeutics Graduate Program, University of Minnesota, Minneapolis, Minnesota 55455, USA

## Abstract

The envelope glycoprotein (Env) trimer on the surface of human immunodeficiency virus type I (HIV-1) mediates viral entry into host CD4+ T cells and is the sole target of neutralizing antibodies. Broadly neutralizing antibodies (bnAbs) that target gp120 V3-glycan of HIV-1 Env trimer are potent and block the entry of diverse HIV-1 strains. Most V3-glycan bnAbs interact, to a different extent, with a glycan attached to N332 but Asn at this position is not absolutely conserved or required for HIV-1 entry based on prevalence of N332 in different circulating HIV-1 strains from diverse clades. Here, we studied the effects of amino acid changes at position 332 of HIV-1_AD8_ Envs on HIV-1 sensitivity to antibodies, cold exposure, and soluble CD4. We further investigated how these changes affect Env function and HIV-1 infectivity *in vitro*. Our results suggest robust tolerability of HIV-1 _AD8_ Env N332 to changes with specific changes that resulted in extended exposure of gp120 V3 loop, which is typically concealed in most primary HIV-1 isolates. Viral evolution leading to Asn at position 332 of HIV_AD8_ Envs is supported by the selection advantage of high levels of cell-cell fusion, transmission, and infectivity even though cell surface expression levels are lower than most N332 variants. Thus, tolerance of HIV-1_AD8_ Envs to different amino acids at position 332 provides increased flexibility to respond to changing conditions/environments and to evade the immune system. Modeling studies of the distance between N332 glycan and specific bnAbs was in agreement with N332 glycan dependency on bnAb neutralization. Overall, our studies provide insights into the contribution of specific amino acids at position 332 to Env antigenicity, stability on ice, and conformational states.

## INTRODUCTION

Human immunodeficiency virus type I (HIV-1) continues to be a major global public health concern with an estimated 39 million people who currently live with HIV (PLWH) worldwide, and over 40 million deaths since the beginning of the HIV-1 pandemic. There is currently no cure for HIV-1 infection but increased access to antiretroviral therapy (ART) has led to improved outcome for PLWH, with ∼80% of patients who are treated with ART showing no detectable viremia (https://www.who.int/). Nevertheless, ART targets viral proteins and cannot eradicate the viral reservoir that contains long-lived cells harboring integrated, replication-competent HIV-1 provirus. Integrated HIV-1 provirus in latent cells is presumably transcriptionally silent but can replenish the viral populations upon treatment discontinuation (1, 2). Some studies documented temporarily or sporadically low-level viral replication (3, 4). Thus, there is a real need to develop an effective HIV-1 vaccine and cure strategies in order to significantly change the course of the HIV-1 pandemic.

HIV-1 envelope glycoproteins (Envs) mediate viral entry into host cells and are the sole target of neutralizing antibodies. On the surface of virions, HIV-1 Envs are assembled into trimers of three subunits, each containing gp120 noncovalently associated with gp41, which anchors the Env trimer to the viral membrane (5). Env interactions with the CD4 receptor lead to structural rearrangements, which reposition the V1/V2 and V3 variable loops of gp120. Movements of these elements outwards lead to Env transitions to an open conformation that exposes the gp120 co-receptor binding region and exposes/forms the gp41 heptad repeat 1 coil on Env surface (5–7). Subsequent binding to CCR5 or CXCR4 coreceptor facilitates further steps during viral entry that culminate in gp41-mediated fusion of the viral and cellular membranes (8–10).

Broadly neutralizing antibodies (bnAbs) against HIV-1 target highly conserved regions of HIV-1 Envs and block viral entry (11–17). At least 5 sites of Env vulnerability have been identified by binding and neutralization activities of different bnAbs (Fig. 1). Vulnerable Env sites include: 1) the CD4 binding site (CD4bs), 2) the V1/V2 loop at the trimer apex, 3) the V3-glycan 4) the gp120-gp41 interface, and 5) the membrane-proximal external region (MPER) of gp41. Robust elicitation of bnAbs by vaccination has been extremely challenging but significant progress in understanding the requirements and/or route for such an immune response has been made over the years (18–23). Nevertheless, despite broad anti-viral activity of bnAbs, HIV-1 Envs evolved diverse mechanisms to escape bnAb neutralization, including conformational dynamics, conformational masking, enhanced cell-to-cell transmission, robustness - which allows HIV-1 Envs to tolerate amino acid changes while maintaining viral entry compatibility - and extensive and alternative glycosylation patterns that protect the underlying protein surface from antibody recognition (24–33).

**Figure 1.**
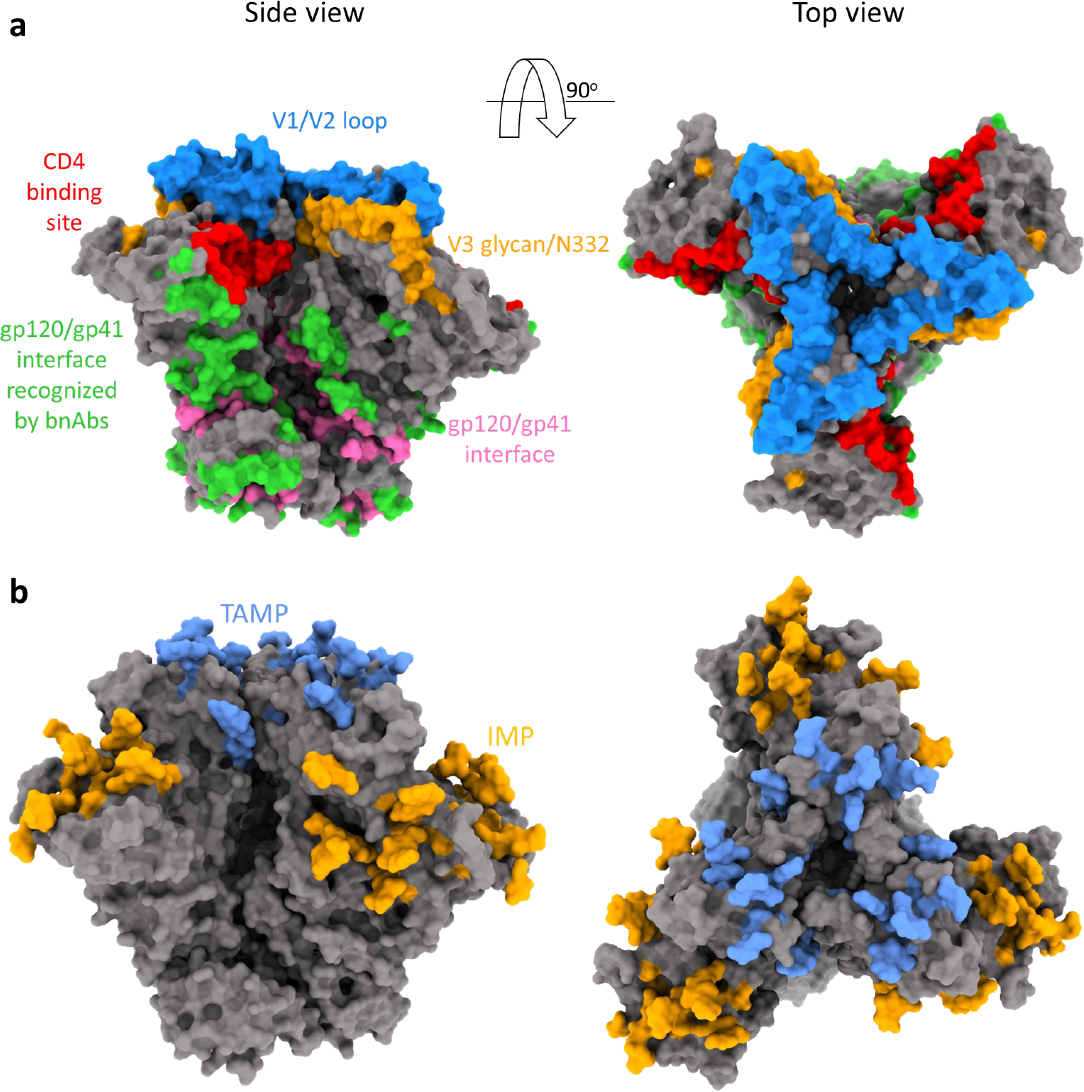
HIV-1 Env vulnerability and dominant glycosylation sites. **a**, Different sites of Env vulnerability defined by bnAb-Env interactions. The gp120 V1/V2 loop, gp120 V3 glycan, and gp120/gp41 interface sites were identified by HIResist (https://hiresist.umn.edu/) using data sourced from CATNAP (Los Alamos National Laboratory). Residues that form the CD4 binding site were labeled according to Kwong et. al., Nature 1998. gp120/gp41 interface was identified by the “protein-protein interface” residue selection tool of Maestro (Schrodinger Suite v21). **b**, HIV-1 Env trimer contains two patches of high mannose, under-processed glycans: Trimer Associated Mannose Patch (TAMP; light blue) and Intrinsic Mannose Patch (IMP; yellow-orange). Glycans on TAMP are added when three gp120 protomers are associated (Moyo et. al., Expert Opin Ther Targets 2021), whereas IMP glycans are located on the gp120 outer domain and can be added at any stage. The TAMP and IMP regions were labeled according to Behrens et. al., 2019. Both the structures were generated using Chimera X and PDB: 8FAD.

Glycans are typically added to eukaryotic proteins in an ordered pattern that involves co-translational addition of a lipid-linked glycosylated oligomannose precursor, Glc_3_Man_9_GlcNAc_2_, to asparagine residues at potential N-linked glycosylation sites (PNGs). These sites are defined by the Asn-X-Ser/Thr sequon, where “X” can be any amino acid except proline (34). After further trimming and processing, eukaryotic glycoproteins may have three general N-glycan types: oligomannose, complex, and hybrid. HIV-1 Envs contain about 75 PNGs and high density of glycans accounts for ∼50% of the trimer mass (34, 35). The high density of PNGs leads to high proportions of glycans that are under processed resulting in high mannose Man_5-9_GlcNAc_2_ glycans that are concentrated in two main regions of HIV-1 Envs: Trimer Associated Mannose Patch (TAMP), and an Intrinsic Mannose Patch (IMP) (Fig. 1) (35, 36). TAMP contains residues N156 and N160 and is formed at the trimer apex when the three gp120 protomers are associated. In contrast, the IMP is primarily mapped to gp120 outer domain and contains the N332 supersite, which is conserved in 73% of ∼30,000 cross-clade Env sequences (HIResist database: https://hiresist.umn.edu/; Milind *et al*. manuscript under review). Although a typical antibody response of humans to foreign antigens is mainly directed against proteins, bnAbs against gp120 V3-glycan can be developed in some PLWH after a long period of infection. V3-glycan bnAbs from PLWH target the IMP glycans, with a specific dependency on the glycan at position N332 for most of them. Structural studies suggest that the N332 glycan is relatively rigid and the dense cluster of neighboring glycans forms an antigenically conserved region. Several studies have demonstrated that HIV-1 often evades N332-directed antibodies by introducing an amino acid change at residue 332 and/or at the related glycosylation sequon. Some V3-glycan bnAbs, such as

PGT121, neutralize HIV-1 strains lacking the N332 glycan that contain neighboring glycans (e.g. N334), whereas Envs of some HIV-1 strains that contain the N332 but lack N334 are resistant to PGT121 but sensitive to 10-1074 (37). For Envs of several HIV-1 strains, neutralization by 10-1074 bnAb may be highly dependent on N332 glycans. Thus, V3-glycan bnAbs differ in their ability to recognize the N332 supersite and, notably, polyclonal V3-glycan targeting bnAbs can arise in a single individual and antibodies from different donors may utilize different modes of recognition (38).

Here, we used the well-studied and highly expressed HIV-1_AD8_ Envs to investigate how changes in residue 332 affect viral fitness, infectivity, and Env function using a variety of assays. We replaced the N332 with all possible 19 amino acids and studied the sensitivity of these variants to bnAbs, ligands preferring open Env conformation such as soluble CD4, and cold, as well as their fusion capacity, and infectivity. Our results provide insights into the relationship between Env function, viral fitness cost, and resistance to V3-glycan bnAbs.

## RESULTS & DISCUSSION

We analyzed the amino acid distribution at position 332 of HIV-1 Env from available viral strains of different clades (Fig. 2) and identified Asn as the most frequent amino acid at this position for most but not all clades. Amino acids other than Asn at position 332 were frequently found in HIV-1 Envs from strains of clade AE that contained mostly the Glu amino acid at this position, whereas stains of clade F2 contained either Thr or Asn at comparable frequencies at this position. Except for clade A1 strains, all clades contained at least 14% strains with an amino acid other than Asn at position 332, suggesting that HIV-1 Envs can tolerate, to some extent, different changes at this site. Thus, to study the contribution of N332 glycan to viral activities, fitness, and resistance to bnAbs targeting the V3-glycan, we introduced amino acid changes in HIV-1_AD8_ Envs to substitute Asn 332 with alternative amino acids. We chose the HIV-1_AD8_ Envs for this study since these Envs are highly expressed, have been extensively investigated in multiple studies, and are typically used to infect non-human primates in a model of HIV-1 infection (39, 40). We used the highly sensitive Cf2-Th/CD4^+^CCR5^+^ target cells and detected high infectivity of all HIV-1_AD8_ mutants except for N332P. Notably, among the natural HIV-1 strains that circulate in the viral population, the prevalence of N332P variant was documented in only 0.06% of clade C strains. This observation suggests that N332P change is either not well tolerated or subject to strong immune pressure within the context of the viral population.

**Figure 2.**
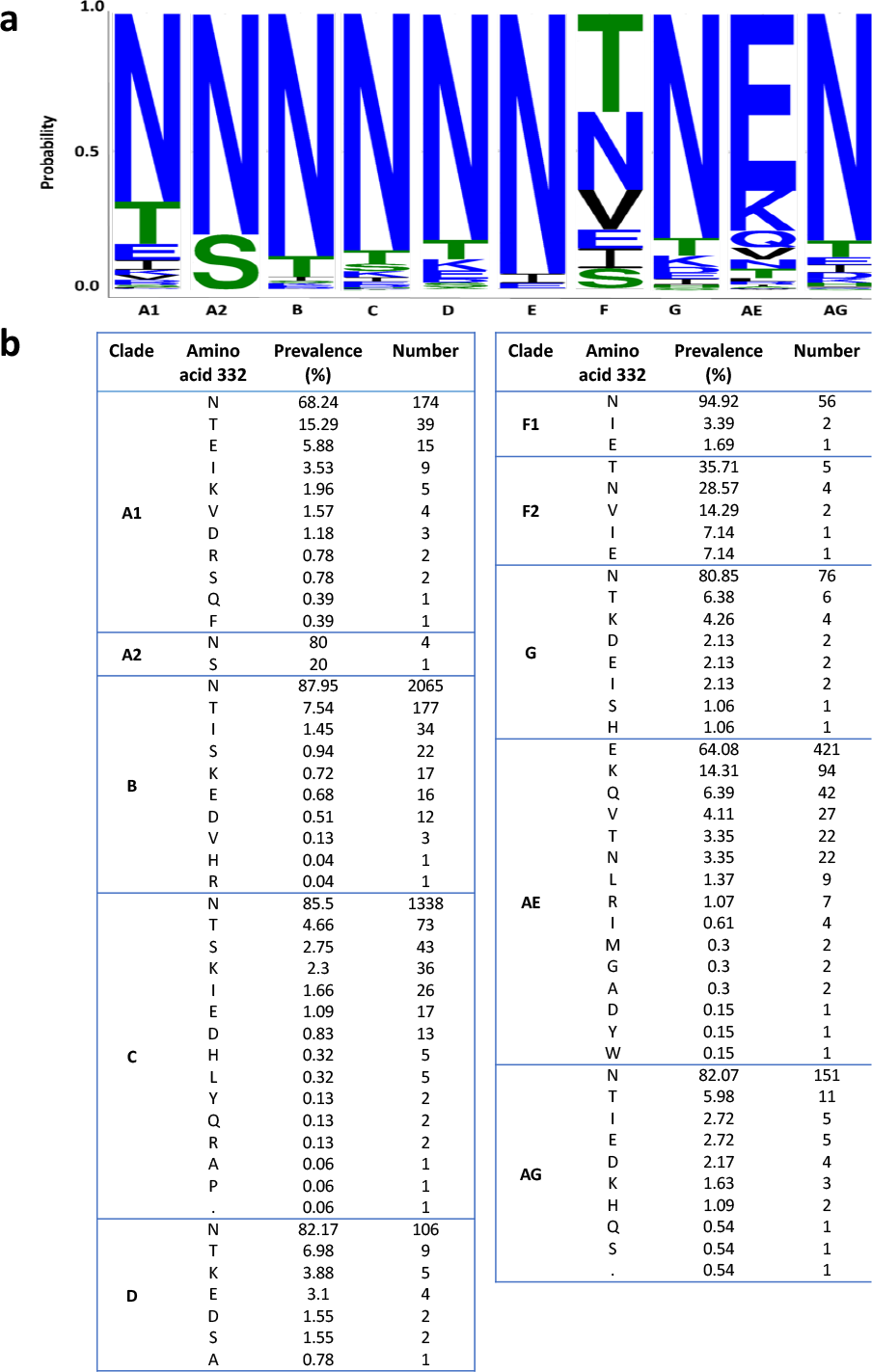
Distribution of amino acids at position 332 of HIV-1 Envs in different HIV-1 strains. **a**, A web logo representation of the prevalence of Asn (N) at position 332 across HIV-1 clades, generated using the AnalyzeAlign (Los Alamos National Laboratory HIV Sequence Database). **b**, Frequency of specific amino acids at position 332 of HIV-1 Envs grouped by different HIV-1 clades.

We evaluated the sensitivity of HIV-1_AD8_ Env variants to the following V3-targeting bnAbs: PGT121, PGT126, PGT128, 10-1074, and 2G12 and compared their sensitivity to the susceptibility of wild type (N332) HIV-1_AD8_ Envs (Figs. 3 and 4). Modeling of the introduced amino acids in the context of HIV-1_AD8_ Envs suggested that none of these amino acids had significant clashes with neighboring residues (Fig. 4a). We detected different patterns of sensitivity of the AD8 Env variants to the V3-glycan bnAbs. PGT bnAbs and 10-1074 exhibited high potency against WT HIV-1_AD8_ Envs with IC_50_ values ranging from 0.01 μg/ml to 0.06 μg/ml and they all were less effective against the AD8 variants, but the level of HIV-1 resistance varied substantially among the different bnAbs. PGT121 and PGT128 still neutralized the AD8 variants but at IC_50_ values that were approximately 2-120-fold higher than the AD8 WT IC_50_ (except for N332C that was completely resistant to PGT128; Fig. 4b and c). In contrast, PGT126 and 10-1074 lost most neutralization activity when tested up to 10 μg/ml against the different AD8 variants. Similarly, 2G12, which only weakly inhibited WT AD8 entry, lost any neutralization activity at or below 10 μg/ml against all AD8 variants. Overall, we detected resistance of all AD8 variants to the V3-glycan bnAbs that varied among the different bnAbs, suggesting that the five V3 targeting glycan bnAbs utilize a diverse mode of binding to the glycan attached to Asn 332. Specifically, 10-1074 and PGT126 rely heavily on the glycan compared to PGT121 and PGT128, which could recognize, to a different extent, the different AD8 variants. Notably, PGT121 exhibited the lowest dependency on Asn at position 332 of HIV-1_AD8_ Envs, maintaining IC_50_ < 1 μg/ml against all HIV-1_AD8_ Env variants and exhibiting the highest resistance to the changes N322L, N332Y, and N332E. This observation is consistent with the potential ability of PGT121 to exhibit promiscuous carbohydrate interactions, recognizing both complex-type N-glycans and high mannose (37). Nevertheless, N332 glycan dependency may vary between diverse HIV-1 strains as the change N332A in HIV-1_JR-CSF_ Envs was reported to abolish any detectable neutralization activity of PGT121 while this change exhibited only 1.4-fold increase in HIV-1_JR-CSF_ resistance to PGT128 (41).

**Figure 3.**
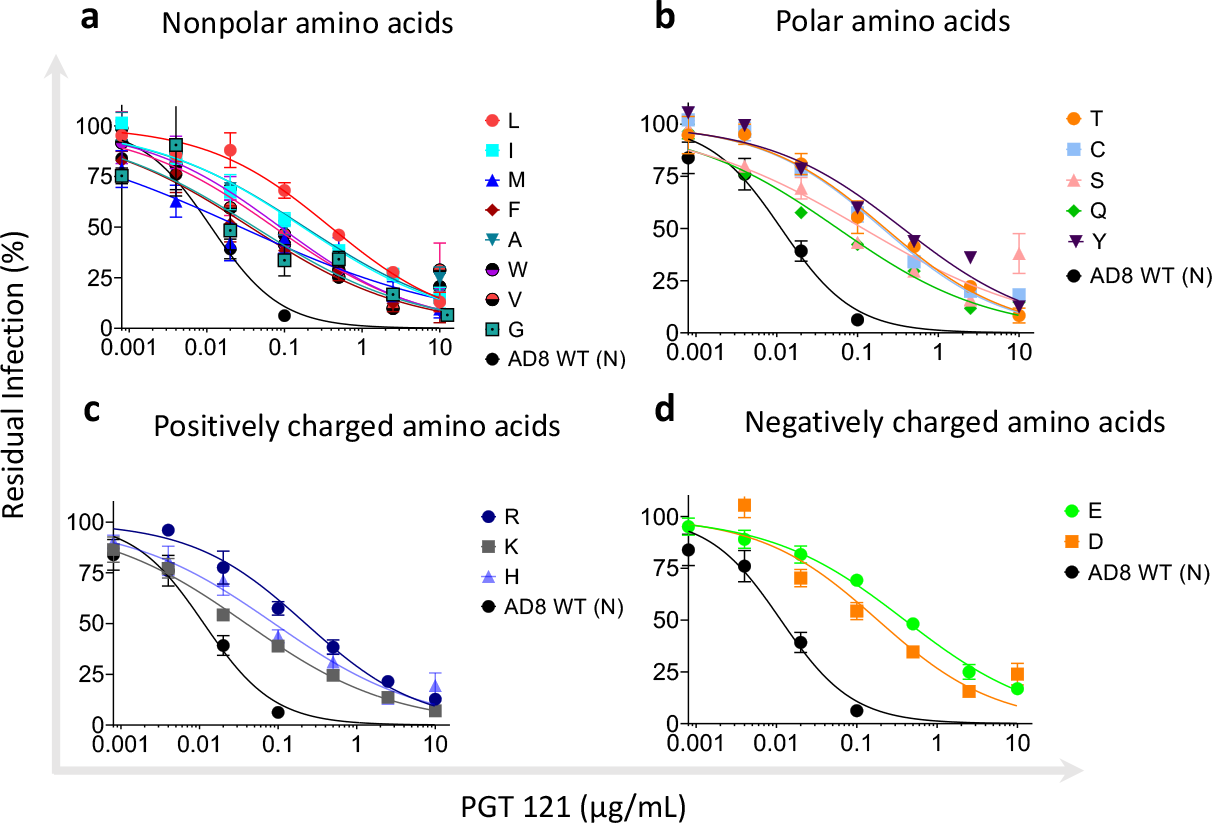
Effects of amino acid variation at residue 332 of HIV-1_AD8_ Env on viral sensitivity to the V3-glycan bnAb PGT121. **a - d**, HIV-1_AD8_ Env variant sensitivity to PGT121. The results were clustered into 4 groups according to the properties of the amino acids with nonpolar (**a**) polar (**b**) positively charges (**c**) and negatively charged (**d**) amino acids. Results are average of at least 2 independent experiments, each performed at least in duplicate.

**Figure 4.**
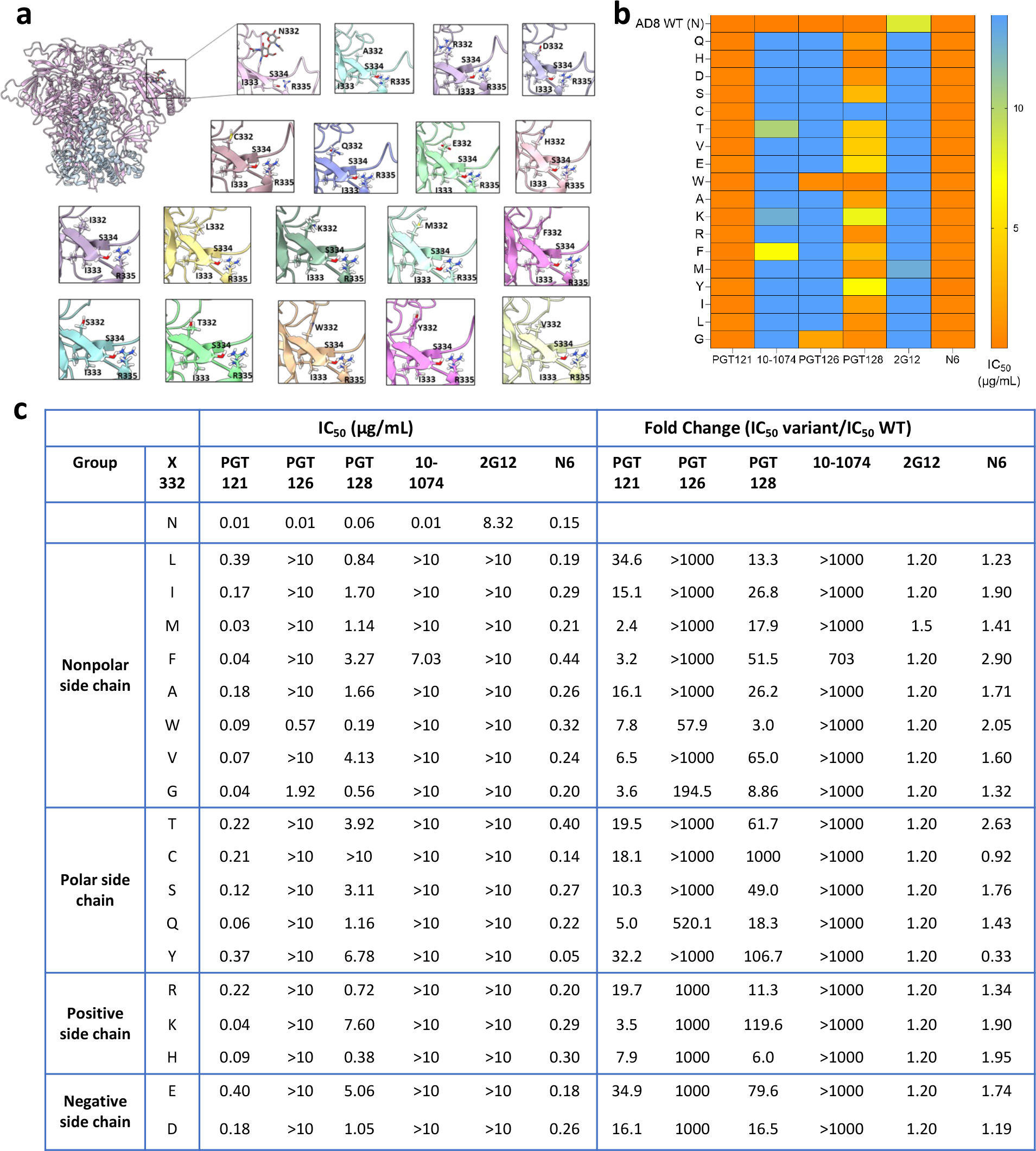
N332 substitutions and patterns of the sensitivity of HIV-1_AD8_ Env variants to V3-glycan bnAbs. **a**, Structures of N332 variants were generated using HIV-1_AD8_ Env trimer structure (PDB: 8FAD) and the mutation tool of Maestro (Schrodinger Suite v21). The structures were then energy-minimized (local minimization) using PRIME-Minimize. **b**, A heat map of HIV-1_AD8_ Env variant sensitivity to several V3-glycan targeting bnAbs was generated based on IC_50_ values (µg/ml) reported in panel c. **c**, IC_50_ values calculated from dose response curves and fold-change of resistance of HIV-1_AD8_ Env N332 variants to each bnAb. Results are average of at least 2 independent experiments, each performed at least in duplicate.

Viruses can infect target cells by diverse pathways and some viruses enter target cells via the endosome where low pH triggers the fusion of viral and cell membranes (42, 43). In contrast, HIV-1 enters target cells by fusion with the plasma membrane at neutral pH, and therefore, HIV-1 Envs evolved to facilitate 2 separate activities: receptor binding and membrane fusion, which are triggered only by receptor/coreceptor binding. Ordered and sequential steps during HIV-1 entry are coordinated by metastability of the Env trimer, which is assembled during synthesis into a ready-to-trigger trimer that adopts a high-energy metastable conformation (stable for a limited time) and stores the energy required to complete the entry process. As a result of Env metastability, some changes in Env residues can exhibit allosteric effects, alter Env conformation and facilitate the exposure of internal Env regions, which are typically concealed in the Envs of most primary strains (6, 44, 45). To evaluate such potential changes, we next studied how different amino acid changes at position 332 affect Env sensitivity to soluble CD4 (sCD4), 19b antibody, which targets an internal epitope of gp120 V3, and exposure to cold (Fig 5). These Env ligands/conditions have been previously associated with Envs that undergo local or global conformational changes and typically exhibit more open Env conformations. Overall, all AD8 variants, as well as AD8 WT, exhibited high sensitivity to sCD4 with IC_50_ values that ranged from 0.01 μM to 0.04 μM. Envs with negatively and positively charged amino acid substitutions at position 332 resulted in slightly more sensitive viruses when they were pseuotyped with these Env variants. As residue 332 is relatively distant from the CD4bs, these observations suggest that Env changes in this residue did not significantly alter the CD4bs (Fig 5). Most amino acid changes at position 332 of Envs resulted in a moderate increase of 19b sensitivity compared with the WT HIV-1_AD8_ Env sensitivity. Unexpectedly, WT HIV-1_AD8_ Envs were sensitive to exposure to cold, which is associated with incompletely closed / partially open / open Env conformations, but all N332 variants except N332G exhibited even higher sensitivity than WT HIV-1_AD8_ Envs to cold exposure. Thus, at least for HIV-1_AD8_ Envs, maintaining the Asn at position 332 was beneficial for limiting the exposure of gp120 V3 loop, which is the target for antibodies developed in PLWH, and for protecting Env elements that are sensitive to cold.

**Figure 5.**
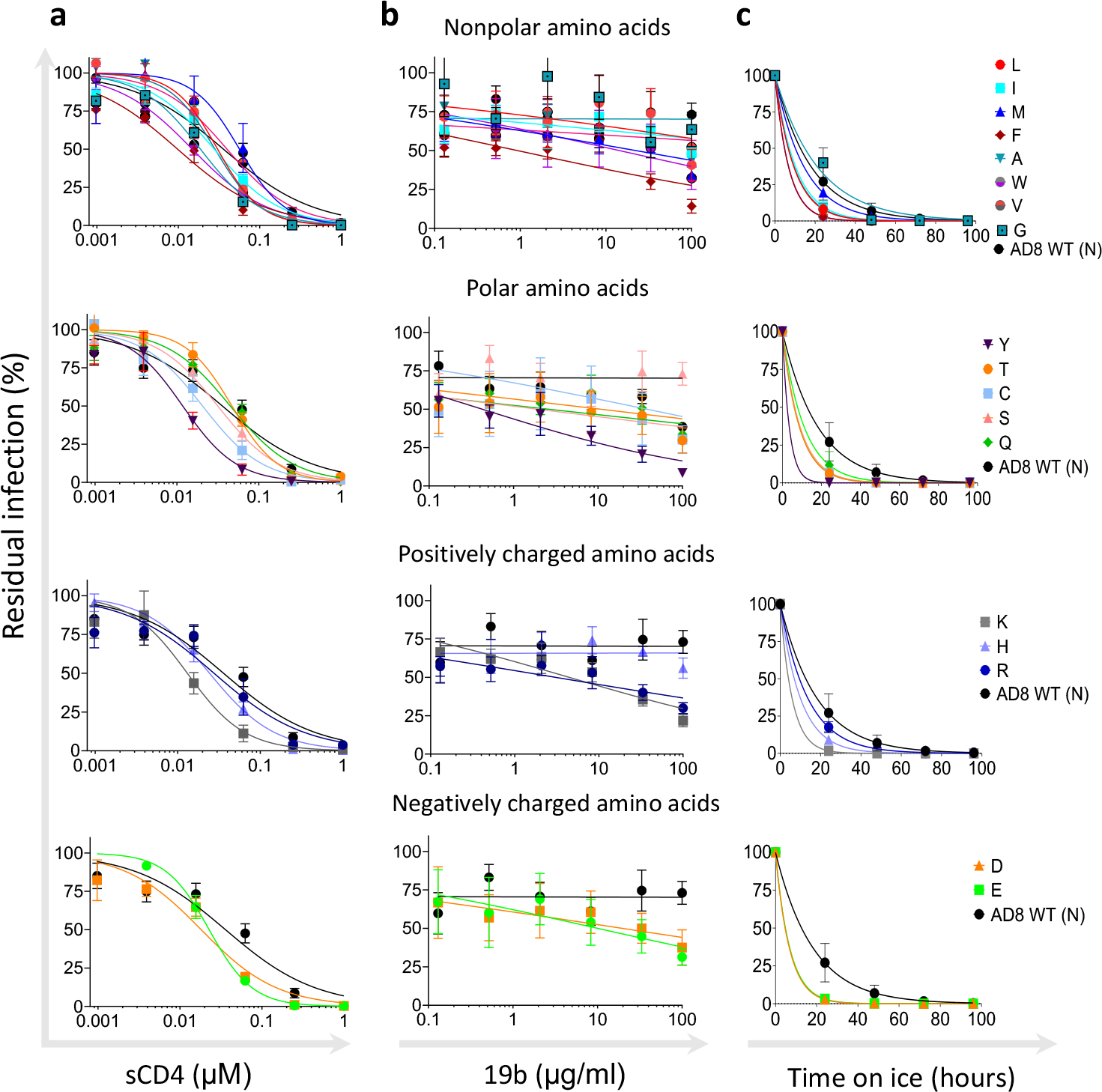
Sensitivity of HIV-1_AD8_ N332 Env variants to ligands and environment changes associated with selectivity toward Env open conformation. Sensitivity of viruses pseudotyped with the indicated Envs to soluble CD4 (sCD4) (**a**); 19b antibody, which targets an internal-epitope of gp120 V3 loop (**b**); and cold exposure (**c**). Results are average of at least 2 independent experiments, each performed at least in duplicate.

To evaluate the effects of amino acid changes at position 332 on HIV-1_AD8_ Env function, we next studied different properties related to the biological activity of HIV-1 Envs (Fig. 6). We assessed the ability of HIV-1 Env variants to mediate cell-cell fusion activity using the well-established HIV-1 Tat reporter system. This assay monitors the fusion of transfected 293T cells coexpressing HIV-1 Tat and Envs with the TZM-bl reporter cells. Except for the N332P, which was poorly infectious, the ability of all other AD8 N332 variants to mediate cell-cell fusion during Amino acid at position 332 of HIV-1_AD8_ Envs a 9-hour incubation was comparable to WT AD8. Nevertheless, the phenotype of the N332G, N332H and N332S variants suggested a slightly delayed kinetics of fusion with low fusion during the first 3 hours of the assay compared to WT AD8. We did not detect significant differences in cell-cell fusion activity between the diverse groups of amino acid substitutions. To investigate the ability of different HIV-1_AD8_ Env variants to mediate the complete entry process, we measured the p24 levels of pseudoviruses displaying the different Env variants, which represents the number of viral particles in the preparation and compared the infectivity of each Env variant at defined p24 levels (Fig. 6). All variants except for the N332P showed high levels of Cf2-Th/CD4^+^CCR5^+^ infection that were comparable to the WT HIV-1_AD8_ Envs. Overall, no significant differences were observed between the diverse groups of amino acids, but the polar amino acid substitutions were slightly less infectious for a defined number of virions (according to p24 levels). As a result, the values of viral infectivity at defined p24 levels of all polar amino acid substitutions were shifted and associated with higher p24 levels relative to the WT AD8 Envs (Fig. 6b). We measured the levels of expression of each Env variant on 293T cells using flow cytometry. Unexpectedly, we detected lower levels of WT HIV-1_AD8_ Envs compared to other variants except for N332P and N332G. This pattern was first detected by the N6 bnAb, which is very potent and recognizes different Env conformations (29), and further confirmed by PGT121, which is highly potent and neutralized all AD8 N332x variants (Fig. 4). We have recently developed an ultrasensitive method to detect transmission of HIV-1 between T cells (CEM to SupT1 cells). The assay uses a reporter lentivector that detects viral transmission in the coculture of viral-producing cells and target cells (31). Reporter protein expression is blocked until HIV-1 completes reverse transcription in target cells, allowing to distinguish between viral-producing cells and target cells to which HIV-1 was transmitted. Using this method, we detected overall high levels of cell-cell transmission between T cells of most variants. Notably, the variations between different members of the same amino acid group were highest among the non-polar amino acid substitutions. Within this group, the HIV-1_AD8_ N332F Env variant exhibited 1.9-fold more efficient cell-cell transmission than the HIV-1_AD8_ N332L Env variant. In summary, WT HIV-1_AD8_ Envs were expressed at lower levels than most variants but maintained comparable cell-cell fusion and transmission activities as well as exhibited comparable or higher viral infectivity.

**Figure 6:**
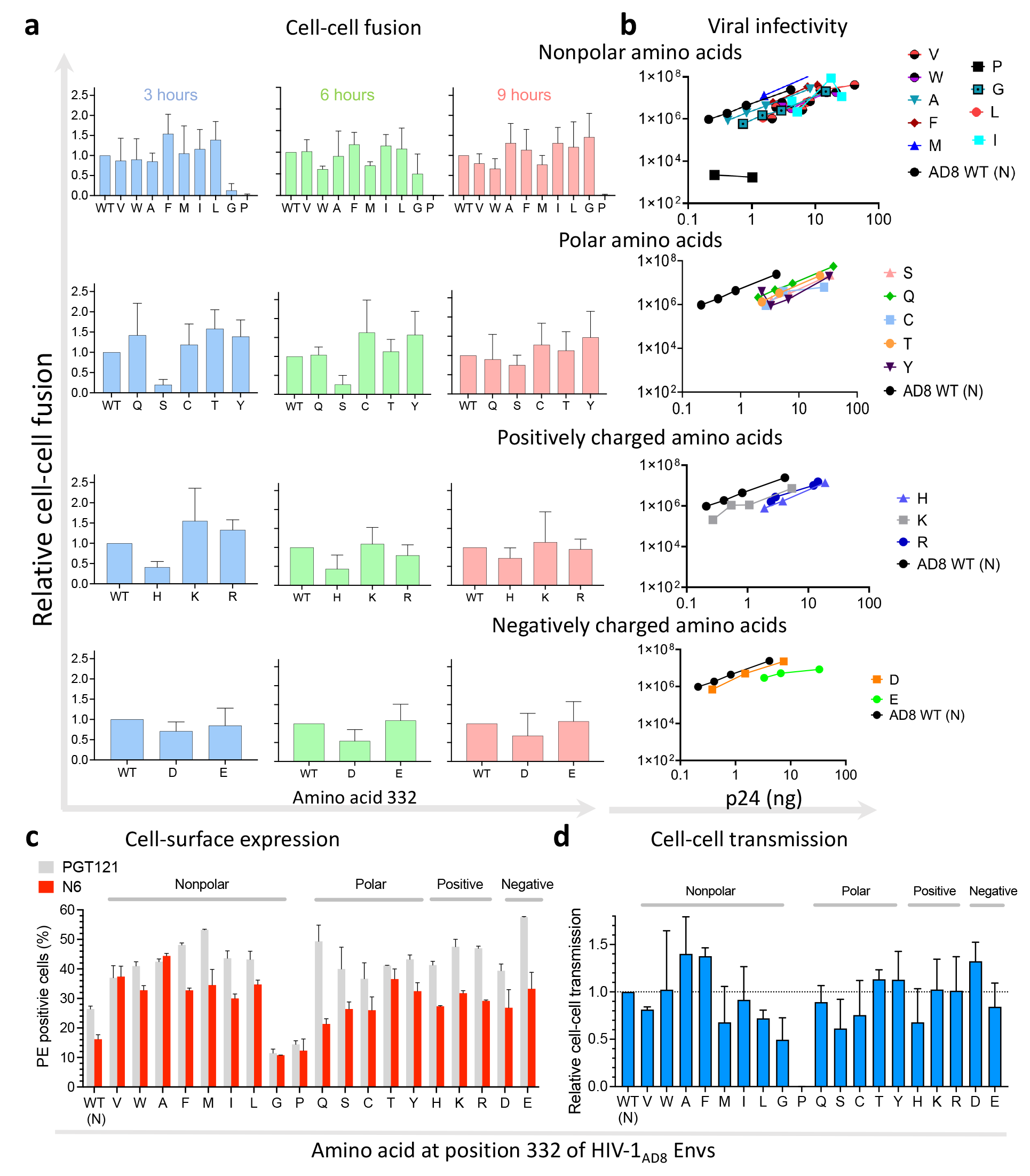
Effects of N332x changes on HIV-1_AD8_ Env function. We measured cell-cell fusion (**a**), assessed infectivity of Envs pseudotyped on HIV-1 particles (**b**), detected Env cell surface expression by flow cytometry using N6 and PGT121 bnAbs (**c**), and evaluated cell-cell transmission (**d**) of HIV-1_AD8_ Env N332x variants. Results are grouped according to the property of the substituted amino acid. Results are representative of at least 2 independent experiments, each performed at least in duplicate.

Correlation analysis of the different parameters that were tested in this study showed a positive correlation between the sensitivity of variants to 1) cold and sCD4, and 2) cold and 19b (Fig. 7). 10-1074 and 2G12 bnAbs exhibited poor neutralization activities (IC_50_ > 10 μg/ml) against most variants that excluded them from the correlation analysis. Notably, sensitivity of the N332x variants to PGT128 bnAb inversely correlated with their sensitivity to cold, suggesting that higher binding affinity to the PGT128 epitope reflects an Env architecture that was better protected from cold.

**Figure 7.**
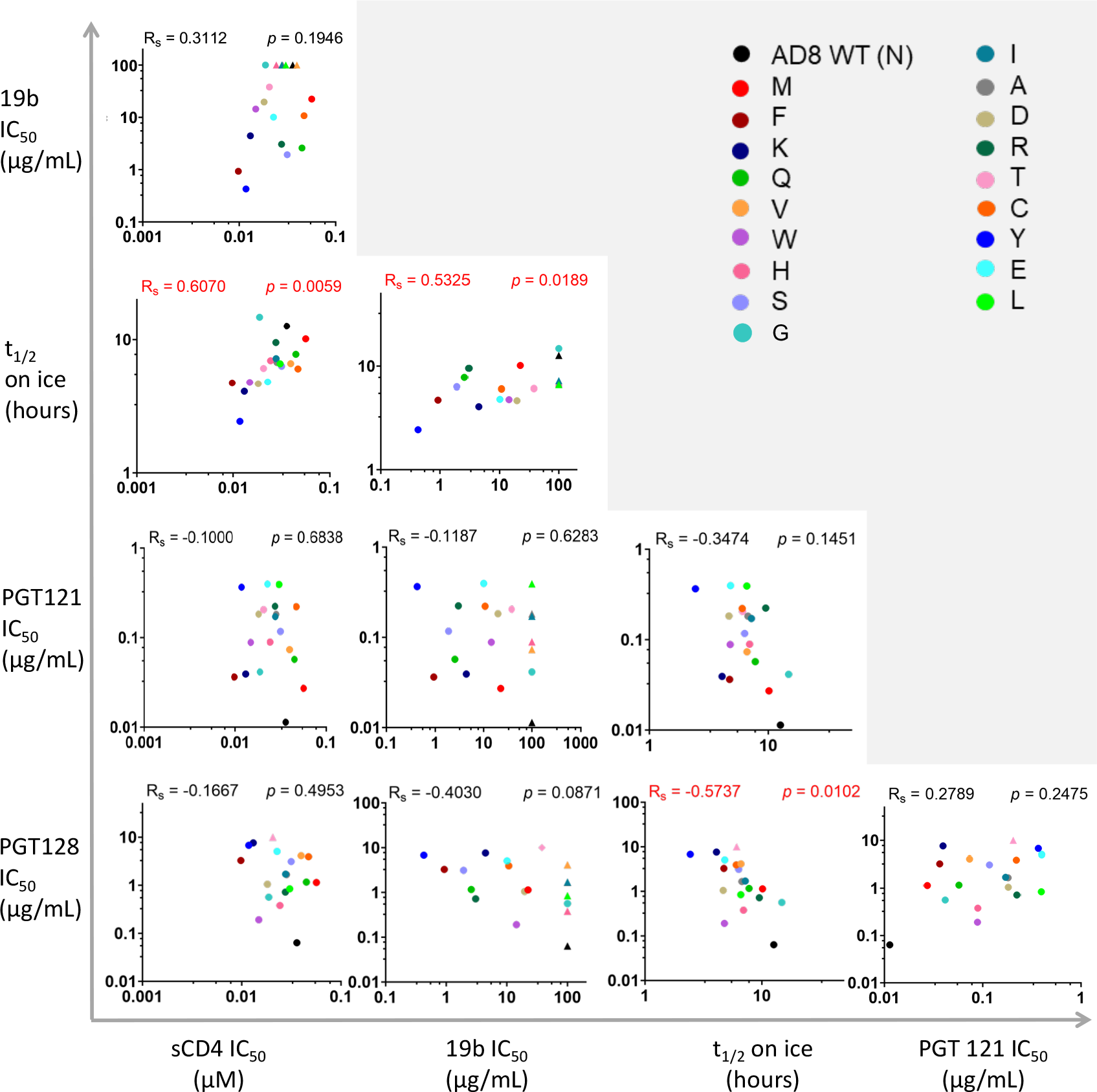
Bi-variable relationship between sensitivities of HIV-1_AD8_ N332x Env variants to multiple ligands and cold exposure. We analyzed the relationship between 5 parameters (IC_50_ of 4 Env ligands and half-life (t_1/2_) on ice) of each N332 variant sensitivity. Rs, non-parametric Spearman correlation coefficient; p, two tailed p value. IC_50_ values are shown as triangles for IC_50_ > 10 µg/mL and as circles for explicit values.

Here we investigated the diverse effects of amino acid changes at position 332 on the sensitivity of HIV-1_AD8_ Envs to different ligands and cold as well as on the ability of N332x variants of HIV-1_AD8_ Env to mediate viral entry, transmit between T cells, and infect target cells. Most amino acid changes were well-tolerated and the related HIV_AD8_ mutants exhibited high levels of expression, high cell-cell fusion, and high infectivity *in vitro*. Viral evolution of Asn at position 332 of HIV_AD8_ Envs is supported by the advantage of high levels of cell-cell fusion, transmission and infectivity even when the cell surface expression levels were lower than most N332x variants. Moreover, most N332x changes increased Env sensitivity to 19b and exposure to cold, suggesting that N332 WT could protect HIV-1 Env from some ligands and/or extreme environmental conditions. Nevertheless, our study points to high robustness of Env function associated with most of the N332x changes. Thus, tolerance of HIV-1_AD8_ Envs to different amino acids at position 332 can provide increased flexibility to respond to changing conditions/environments.

The different pattern of sensitivity to the V3-glycan bnAbs indicated diverse dependency in the N332 glycan and potentially different antibody approach to the V3-glycan epitope. To obtain additional insights into the binding mode of the different V3-glycan bnAbs to their epitope, we modeled bnAb binding using an available cryo-EM structure of the full-length HIV-1_AD8_ Envs (46). We superimposed HIV-1_AD8_ Env structure on available structures of HIV-1_BG505_ SOSIP.664 in complex with the different V3-glycan bnAbs and estimated the distance between these bnAbs and HIV-1_AD8_ Env N332 glycan (Fig. 8). Distances between PGT121, PGT128, and 10-1074 and N332 glycan were consistent with our results of bnAb neutralization dependency on N332 glycan (Figs. 3 and 4). PGT121 showed the lowest dependency and effectively neutralized all the N332 variants (although these variants were less efficiently neutralized than WT N332) and it was also the most distant from N332 glycan among the three bnAbs in this model. In contrast, 10-1074 showed poor neutralization activity in the absence of N332 glycan and it was positioned in close proximity to the N332 glycan in our model (Fig. 8). Of note, dose-response curve of HIV_AD8_ WT to increasing 10-1074 concentrations showed complete neutralization at high concentration of 10-1074, suggesting that the addition of N332 glycan is homogenous among all pseudoviruses produced in 293T cells. HIV_AD8_ WT was relatively resistant to the 2G12 antibody, and most variants were completely resistant to this antibody up to 10 μg/ml; this observation complicated any conclusion for the distance dependency of 2G12.

**Figure 8.**
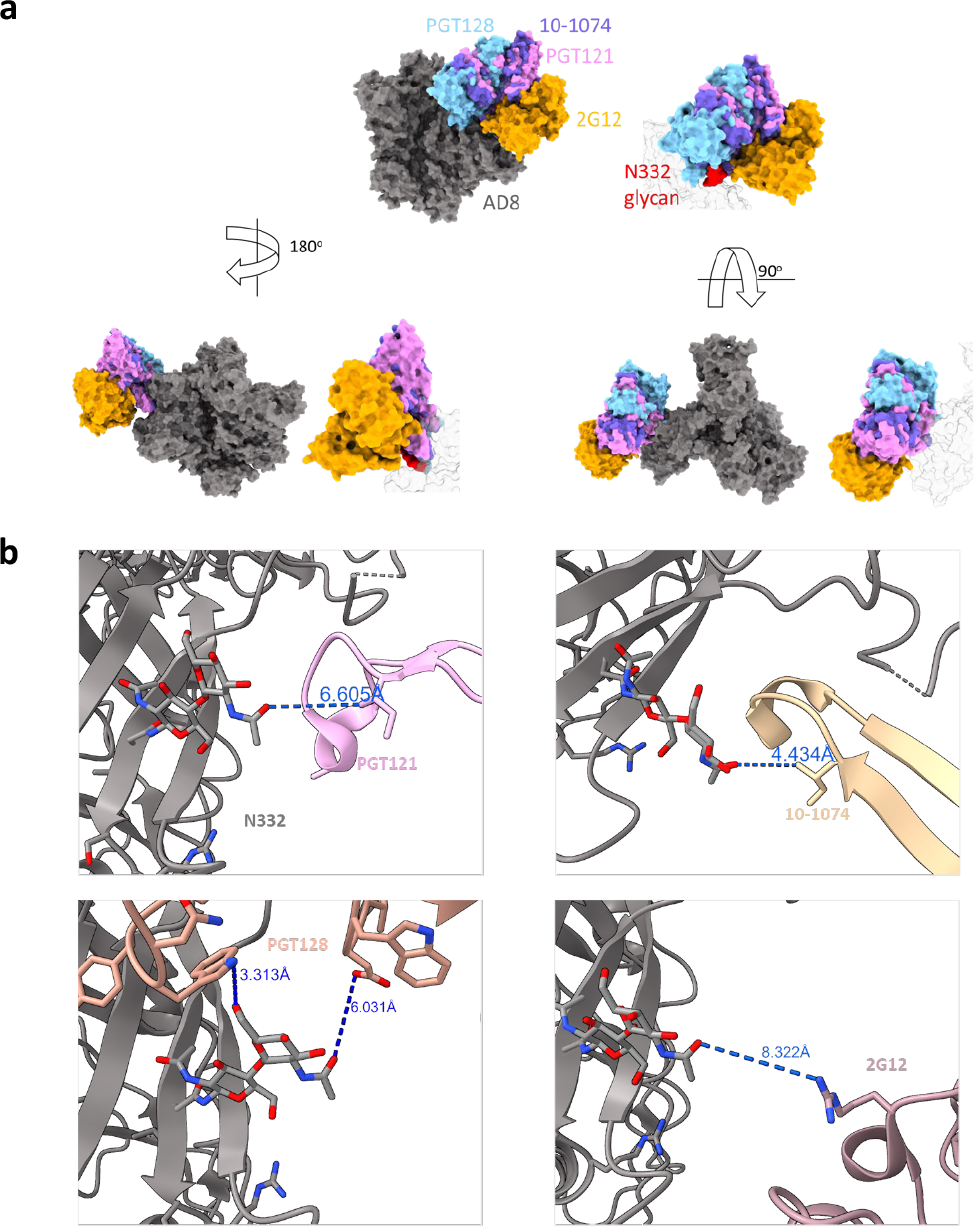
Molecular modeling of potential interactions of different V3-glycan targeting bnAbs with HIV-1_AD8_ Envs. **a**, We used available structures of different V3-glycan bnAbs in complex with BG505 SOSIP.665 (PDBs: 5ACO, 6OCZ, 6UDJ, and 7UOJ) to evaluate of the interactions of these bnAbs with the IMP region of HIV-1_AD8_ Env. We superimposed HIV-1_AD8_ Env structure (PDB 8FAD) with the bnAb-BG505 SOSIP complex structures using using Chimera X matchmaker tool and then removed the BG505 SOSIP structure. 2G12 complex included 2 Fab and one of these was removed for clarity. b, Modeling the HIV-1_AD8_ Env and N322 interactions with V3-glycan bnAbs. The models show the closest distance between each bnAb and N332 glycan of HIV-1_AD8_ Envs (PDB 8FAD).

The prevalence of HIV-1 strains lacking N332 among circulating strains of HIV-1 suggests that the glycan at this position is not absolutely necessary for HIV-1 Env function or for HIV-1 replication. Of note, some transmitted / founder HIV-1 strains, which can establish HIV-1 infection in vivo, lack N332 glycan and have been previously identified and isolated (e.g., BG505). Nevertheless, most HIV-1 strains do contain Asn at position 332. One potential explanation for this pattern of distribution may be temporal and alternative adaptability of HIV-1 Env to V3-glycan bnAbs and antibodies typically present in patient serum (Kshitij Wagh personal communication). Thus, switching between different amino acids can confer resistance against different types of antibodies at different time points during HIV-1 infection. Notably, changes of Asn to some charged amino acids as well as to Tyr and Phe resulted in significant exposure of the gp120 V3 loop of HIV-1_AD8_ Env (clade B) according to the sensitivity of these variants to the internal-epitope antibody 19b, which targets the gp120 V3 loop (Fig. 5). Consistent with this observation, the prevalence of these changes in the viral population of Clade B HIV-1 strains is low (Fig. 2). Our study provides insights into the role of amino acids at position 332 of HIV-1 Envs and the effects of different changes on HIV-1 fitness and sensitivity to antibodies.

## MATERIALS & METHODS

### Plasmid construction

To generate the HIV-1_AD8_ *env* mutants, we first used the primer AD8-M_N332x_F (5’-cgccaggcccactgcnnkatcagccgcaccaag-3’) that contained the NNK degenerate codons at position 332 to create a small library of mutants and screened for single mutants. The library was generated either using QuikChange multisite lightning site directed mutagenesis kit (Agilent) or by PCR amplification followed by Gibson assembly into the original pCDNA-AD8 plasmid and had the diversity of 32 alternative codons, including only one stop codon. Due to biased toward specific mutations, we also designed specific primers for site directed mutagenesis of the following mutants that could not be recovered directly from the library:

N332A

AD8M N332A F: 5’-CCAGGCCCCTGCGCCATCAGCCGCACC-3’ AD8M N332A R: 5’-GGTGCGGCTGATGGCGCAGTGGGCCTGG-3’

N332E

AD8M N332E F: 5’-GCCAGGCCCACTGCGAAATCAGCCGCACCAA-3’ AD8M N332E R: 5’-TTGGTGCGGCTGATTTCGCAGTGGGCCTGGC-3’

N332R

AD8M N332R F: 5’-CCAGGCCCACTGCAGGATCAGCCGCACCAA-3’ AD8M N332R R: 5’-TTGGTGCGGCTGATCCTGCAGTGGGCCTGG-3’

N332T

AD8M N332T F: 5’-CAGGCCCACTGCACCATCAGCCGCACC-3’ AD8M N332T R: 5’-GGTGCGGCTGATGGTGCAGTGGGCCTG-3’

N332V

AD8M N332V F: 5’-CCAGGCCCACTGCGTCATCAGCCGCACC-3’ AD8M N332V R: 5’-GGTGCGGCTGATGACGCAGTGGGCCTGG-3’

N332W

AD8M N332W F: 5’-CGCCAGGCCCACTGCTGGATCAGCCGCACCAAG-3’ AD8M N332W R: 5’-CTTGGTGCGGCTGATCCAGCAGTGGGCCTGGCG-3’

The correct DNA sequences were verified by Sanger sequencing.

### Cell lines

293T cell were purchased from the American Type Culture Collection (ATCC) and TZM-bl cells were obtained from the NIH AIDS Reagent Program. Cf2-Th/CD4^+^CCR5^+^ cells were a kind gift from Joseph Sodroski (Dana-Farber Cancer Institute). CEM CD4+ cells were obtained from the NIH AIDS Reagent Program, and SupT1.CCR5 (SupT1.R5) cells stably expressing the human CCR5 coreceptor were a kind gift from James Hoxie (University of Pennsylvania). All cell lines were tested negative for mycoplasma. 293T cells were grown in Dulbecco’s Modified Eagle Medium (DMEM) with 10% Fetal Bovine Serum (FBS), 10mL of 100μg/mL streptomycin and 100units/mL of penicillin (all from Gibco, ThermoFisher Scientific). Cf2-Th/CD4^+^CCR5^+^ cells grown in the presence of 400 µg/ml G418 and 200 µg/ml hygromycin B selection antibiotics (both from Invitrogen, ThermoFisher Scientific). CEM, SupT1.CCR5 cells were maintained in RPMI 1640 medium containing 10% FBS, 2 mM glutamine, 100 U/ml penicillin, and 100 µg/ml streptomycin.

### Production of single-round pseudoviruses

We produced pseudoviruses as previously described (47–49) using Effectene (Qiagen). Briefly, we co-transfected 293T cells with three plasmids: an envelope expressing plasmid, pHIVec2.luc reporter plasmid and psPAX2 packaging plasmid (catalog number 11348, NIH AIDS Reagent Program) at a ratio of 1:6:3, respectively. After a 48-hour incubation, cell supernatant was collected and centrifuged for 10 minutes at 600-900x g at 4°C. The amount of p24 in the supernatant was measured using HIV-1 p24 ELISA Kit (cat# XB-1000, Xpress Bio) and pseudovirus-containing supernatant was frozen in single-use aliquots at -80°C.

### Viral infection assay

Viral infection assay was performed as previously described (25, 44, 48, 50–52). HIV-1 Env ligands (antibodies and sCD4) were serially diluted in DMEM, and 30μL of each tested concentration was dispensed into a single well (each concentration was tested in duplicate in each experiment) of a 96 well-white plate (Greiner Bio-One, NC). Pseudoviruses were thawed at 37°C for 1.5 min and 30μL were added to each well. After a brief incubation, 30μL of 2.0 x10^5^ Cf2-Th/CD4^+^CCR5^+^ target cells/mL suspended in DMEM were added to each well. Pseudoviruses were typically titered on Cf2-Th/CD4^+^CCR5^+^ target and volumes corresponding to ∼1-3 million relative light units (2-sec integration using Centro LB 960 or Centro XS^3^ LB 960 luminometer) were used in all experiments. After 48-hour incubation, the medium was aspirated, and the cells were lysed with 30μL of Lysis Buffer (25mM Tris, Trans-1,2-Diaminocyclohexane-N,N,N’,N’ tetra acetic acid monohydrate, 1% triton, 10% glycerol, and 2mM Dithiothreitol, with pH adjusted to 7.8). Firefly luciferase activity was measured with 2-sec integration time using a Centro LB 960 (or Centro XS^3^ LB 960) luminometer (Berthold Technologies, TN, USA).

To determine the effect of cold exposure on viral infectivity, single-use aliquots of the pseudoviruses were thawed at 24-hour intervals (day 0, 1,2,3, and 4 of the experiment) at 37°C for 1.5 min and then incubated on ice for a specified time (24-96 hours). At the end of the incubation period, equal amounts of pseudoviruses were added to Cf2-Th/CD4^+^CCR5^+^ target cells and infectivity was measured after 48 hours as described above.

Dose-response curves were generated by fitting the data to a four-parameter logistic equation using Prism 9 program (GraphPad, San Diego, CA), which calculated IC_50_ and SEM according to the fitted curves (53, 54). Dose-response curves of viral infectivity decay during cold exposure was fitted to one-phase exponential decay using the Prism 9 Program. Statistical significance (Fig. 7) was tested using nonparametric Spearman analysis included in the Prism 9 Program and correlation values along with two-tailed p values are reported (Fig. 7). The number of experiments repeated, and replicates are provided in the Figure Legends.

### Cell-cell fusion assay

Cell-cell fusion was measured as previously described with HIV-1 Tat as a transactivator and TZM-bl cells as reporter target cells (48, 55). In a 6-well plate, 5.0 x10^5^ 293T cells were co-transfected with HIV-1 Env-expressing and HIV-1 Tat-expressing plasmids at a ratio of 1:6 using Effectene (Qiagen) and incubated for 48 hours. After 24 hours, 10,000 TZM-bl reporter cells/well were added to 96-well white plates (Greiner Bio-One, NC) and the plates were incubated overnight. At the end of the 48-hour incubation, the 293T cells were detached using 5mM EDTA/PBS, washed once, resuspended in fresh media and 10,000 cells were added to 96 plates containing the pre-incubated TZM-bl cells. Plates were incubated for 3, 6, and 9 hours, and cells were then lysed; firefly luciferase activity was measured with 2-sec integration time using a Centro XS^3^ LB 960 luminometer (Berthold Technologies, TN, USA).

### Cell-to-cell transmission assay

Cell-cell transmission was measured as previously described (31). Briefly, 10^6^ CEM CD4+ T cells were washed once with PBS (in 1.5 ml tubes), and electroporated in buffer R (Neon transfection reagent; Invitrogen, USA) with 3 µg of the reporter plasmid pUCHR-EF1a-inNluc, 2 µg of the packaging plasmid pNL4-3ΔEnv, and 1 µg of AD8 Env expression plasmids (WT or N332x variants). Cells were electroporated with Neon Transfection System (Invitrogen, USA) using 100 µl tips with the following setting: 1230 V, 40 ms x 1 pulse. Electroporated cells were added to 1 ml of culture medium containing 5x10^5^ SupT1.CCR5 target cells. Cell mixture was washed 2 times with culture medium to remove plasmid DNA and incubated in a final volume of 1.5 ml growth medium in a 12-well plate for 72 hours. Efficiency of cell-cell transmission was evaluated by measuring nano luciferase (Nluc) activity as follows. Samples were transferred from wells of the 12-well plate to 1.5 mL tubes, cells were centrifuged at 1,000 x g for 3 min and lysed by adding 50 µl of Promega Glo lysis buffer. Cell lysates were clarified by centrifugation at 10,000 x g for 2 min, and transferred to a white 96-well plate. After 3-minute incubation we added 30 µl of NanoGlo Luciferase substrate (Promega) per well and nano luciferase activity was measured by Centro XS^3^ LB 960 luminometer (Berthold Technologies).

### Molecular modeling

Amino acid changes of residue 332 of HIV-1_AD8_ Envs (Fig. 4a) were generated using available HIV-1_AD8_ Env structure (PDB: 8FAD) and the Mutation Tool in Maestro graphical user interface (Schrödinger Suite v21; Schrödinger, LLC, New York, NY). The structures were further locally minimized (only the potential N-glycosylation sequon (residues X332-R335) was minimized) using the PRIME-Minimize Panel. Default solvation model VSGB was applied with an OPLS4 force field, and RMS gradient for convergence was 0.01 kcal/mol/Å. Structure modeling (Figs. 7 and 8) was performed using Maestro graphical user interface (Schrodinger Suite v21), UCSF Chimera X (56), HIV-1_AD8_ Env trimer structure (PDB: 8FAD) and the following structures of bnAbs in complexes with BG505 SOSIP: 1) PGT121 - PDB: 7UOJ, 2) PGT128 - PDB: 5ACO, 3) 10-1074 - PDB:6UDJ, and 4) 2G12 - PDB: 6OZC. We used the HIV-1_AD8_ Env trimer structure (PDB: 8FAD) as a template and superimposed bnAb-BG505 SOSIP complex structures on the HIV-1_AD8_ Env trimer structure using ChimeraX Matchmaker Tool. We generated a separate model for each bnAb in complex with HIV-1_AD8_ Env trimer. BG505.664 trimer and the second Fab of 2G12 were then removed from the display. Distances between the closest bnAb residue and AD8 N332 glycan were measured using Chimera X Distance tool.

## AKNOWLEDGMENTS

We thank the NIH AIDS Reagent Program, Division of AIDS, NIAID for providing Env-expressing plasmids and broadly neutralizing antibodies. Molecular graphics and analyses performed with UCSF ChimeraX, developed by the Resource for Biocomputing, Visualization, and Informatics at the University of California, San Francisco, with support from National Institutes of Health R01-GM129325 and the Office of Cyber Infrastructure and Computational Biology, National Institute of Allergy and Infectious Diseases.

## DATA AVAILABILITY

Data is available upon request.

## REFERENCES

1. Bruner KM, Wang Z, Simonetti FR, Bender AM, Kwon KJ, Sengupta S, Fray EJ, Beg SA, Antar AAR, Jenike KM, Bertagnolli LN, Capoferri AA, Kufera JT, Timmons A, Nobles C, Gregg J, Wada N, Ho YC, Zhang H, Margolick JB, Blankson JN, Deeks SG, Bushman FD, Siliciano JD, Laird GM, Siliciano RF. 2019. A quantitative approach for measuring the reservoir of latent HIV-1 proviruses. Nature 566:120–125.

2. Chun TW, Carruth L, Finzi D, Shen X, DiGiuseppe JA, Taylor H, Hermankova M, Chadwick K, Margolick J, Quinn TC, Kuo YH, Brookmeyer R, Zeiger MA, Barditch-Crovo P, Siliciano RF. 1997. Quantification of latent tissue reservoirs and total body viral load in HIV-1 infection. Nature 387:183–188.

3. Wietgrefe SW, Duan L, Anderson J, Marqués G, Sanders M, Cummins NW, Badley AD, Dobrowolski C, Karn J, Pagliuzza A, Chomont N, Sannier G, Dubé M, Kaufmann DE, Zuck P, Wu G, Howell BJ, Reilly C, Herschhorn A, Schacker TW, Haase AT. 2022. Detecting Sources of Immune Activation and Viral Rebound in HIV Infection. J Virol 96:e0088522.

4. Boritz EA, Darko S, Swaszek L, Wolf G, Wells D, Wu X, Henry AR, Laboune F, Hu J, Ambrozak D, Hughes MS, Hoh R, Casazza JP, Vostal A, Bunis D, Nganou-Makamdop K, Lee JS, Migueles SA, Koup RA, Connors M, Moir S, Schacker T, Maldarelli F, Hughes SH, Deeks SG, Douek DC. 2016. Multiple Origins of Virus Persistence during Natural Control of HIV Infection. Cell 166:1004–1015.

5. Liu J, Bartesaghi A, Borgnia MJ, Sapiro G, Subramaniam S. 2008. Molecular architecture of native HIV-1 gp120 trimers. Nature 455:109–13.

6. Herschhorn A, Gu C, Espy N, Richard J, Finzi A, Sodroski JG. 2014. A broad HIV-1 inhibitor blocks envelope glycoprotein transitions critical for entry. Nature Chemical Biology 10:845–852.

7. Si Z, Madani N, Cox JM, Chruma JJ, Klein JC, Schön A, Phan N, Wang L, Biorn AC, Cocklin S, Chaiken I, Freire E, Smith AB, Sodroski JG. 2004. Small-molecule inhibitors of HIV-1 entry block receptor-induced conformational changes in the viral envelope glycoproteins. Proceedings of the National Academy of Sciences of the United States of America 101:5036–41.

8. Weissenhorn W, Dessen A, Harrison SC, Skehel JJ, Wiley DC. 1997. Atomic structure of the ectodomain from HIV-1 gp41. Nature 387:426–430.

9. Lu M, Blacklow SC, Kim PS. 1995. A trimeric structural domain of the HIV-1 transmembrane glycoprotein. Nature structural biology 2:1075–82.

10. Melikyan GB, Markosyan RM, Hemmati H, Delmedico MK, Lambert DM, Cohen FS. 2000. Evidence that the transition of HIV-1 gp41 into a six-helix bundle, not the bundle configuration, induces membrane fusion. The Journal of cell biology 151:413–23.

11. Wu X, Yang Z-Y, Li Y, Hogerkorp C-M, Schief WR, Seaman MS, Zhou T, Schmidt SD, Wu L, Xu L, Longo NS, McKee K, O’Dell S, Louder MK, Wycuff DL, Feng Y, Nason M, Doria-Rose N, Connors M, Kwong PD, Roederer M, Wyatt RT, Nabel GJ, Mascola JR. 2010. Rational design of envelope identifies broadly neutralizing human monoclonal antibodies to HIV-1. 5993. Science (New York, NY) 329:856–61.

12. Zhou T, Georgiev I, Wu X, Yang ZY, Dai K, Finzi A, Kwon YD, Scheid JF, Shi W, Xu L, Yang Y, Zhu J, Nussenzweig MC, Sodroski J, Shapiro L, Nabel GJ, Mascola JR, Kwong PD. 2010. Structural basis for broad and potent neutralization of HIV-1 by antibody VRC01. Science 329:811–817.

13. Burton DR, Pyati J, Koduri R, Sharp SJ, Thornton GB, Parren PW, Sawyer LS, Hendry RM, Dunlop N, Nara PL. 1994. Efficient neutralization of primary isolates of HIV-1 by a recombinant human monoclonal antibody. Science (New York, NY) 266:1024–7.

14. Caskey M, Klein F, Lorenzi JCC, Seaman MS, West AP, Buckley N, Kremer G, Nogueira L, Braunschweig M, Scheid JF, Horwitz JA, Shimeliovich I, Ben-Avraham S, Witmer-Pack M, Platten M, Lehmann C, Burke LA, Hawthorne T, Gorelick RJ, Walker BD, Keler T, Gulick RM, Fätkenheuer G, Schlesinger SJ, Nussenzweig MC. 2015. Viraemia suppressed in HIV-1-infected humans by broadly neutralizing antibody 3BNC117. Nature 522:487–491.

15. Huang J, Ofek G, Laub L, Louder MK, Doria-Rose NA, Longo NS, Imamichi H, Bailer RT, Chakrabarti B, Sharma SK, Alam SM, Wang T, Yang Y, Zhang B, Migueles SA, Wyatt R, Haynes BF, Kwong PD, Mascola JR, Connors M. 2012. Broad and potent neutralization of HIV-1 by a gp41-specific human antibody. Nature 491:406–12.

16. Huang J, Kang BH, Ishida E, Zhou T, Griesman T, Sheng Z, Wu F, Doria-Rose NA, Zhang B, McKee K, O’Dell S, Chuang G-Y, Druz A, Georgiev IS, Schramm CA, Zheng A, Joyce MG, Asokan M, Ransier A, Darko S, Migueles SA, Bailer RT, Louder MK, Alam SM, Parks R, Kelsoe G, Von Holle T, Haynes BF, Douek DC, Hirsch V, Seaman MS, Shapiro L, Mascola JR, Kwong PD, Connors M. 2016. Identification of a CD4-Binding-Site Antibody to HIV that Evolved Near-Pan Neutralization Breadth. Immunity 45:1108–1121.

17. Julien J-P, Sok D, Khayat R, Lee JH, Doores KJ, Walker LM, Ramos A, Diwanji DC, Pejchal R, Cupo A, Katpally U, Depetris RS, Stanfield RL, McBride R, Marozsan AJ, Paulson JC, Sanders RW, Moore JP, Burton DR, Poignard P, Ward AB, Wilson IA. 2013. Broadly Neutralizing Antibody PGT121 Allosterically Modulates CD4 Binding via Recognition of the HIV-1 gp120 V3 Base and Multiple Surrounding Glycans. PLoS Pathogens 9:e1003342.

18. Ratnapriya S, Perez-Greene E, Schifanella L, Herschhorn A. 2022. Adjuvant-mediated enhancement of the immune response to HIV vaccines. FEBS J 289:3317–3334.

19. Abbott RK, Lee JH, Menis S, Skog P, Rossi M, Ota T, Kulp DW, Bhullar D, Kalyuzhniy O, Havenar-Daughton C, Schief WR, Nemazee D, Crotty S. 2018. Precursor Frequency and Affinity Determine B Cell Competitive Fitness in Germinal Centers, Tested with Germline-Targeting HIV Vaccine Immunogens. Immunity 48:133–146.e6.

20. Ahlers JD. 2014. All eyes on the next generation of HIV vaccines: Strategies for inducing a broadly neutralizing antibody response. Discovery Medicine.

21. Kwong PD, Mascola JR. 2018. HIV-1 Vaccines Based on Antibody Identification, B Cell Ontogeny, and Epitope Structure. Immunity 48:855–871.

22. Andrabi R, Voss JE, Liang CH, Briney B, McCoy LE, Wu CY, Wong CH, Poignard P, Burton DR. 2015. Identification of Common Features in Prototype Broadly Neutralizing Antibodies to HIV Envelope V2 Apex to Facilitate Vaccine Design. Immunity 43:959–973.

23. Bricault CA, Yusim K, Seaman MS, Yoon H, Theiler J, Giorgi EE, Wagh K, Theiler M, Hraber P, Macke JP, Kreider EF, Learn GH, Hahn BH, Scheid JF, Kovacs JM, Shields JL, Lavine CL, Ghantous F, Rist M, Bayne MG, Neubauer GH, McMahan K, Peng H, Chéneau C, Jones JJ, Zeng J, Ochsenbauer C, Nkolola JP, Stephenson KE, Chen B, Gnanakaran S, Bonsignori M, Williams LD, Haynes BF, Doria-Rose N, Mascola JR, Montefiori DC, Barouch DH, Korber B. 2019. HIV-1 Neutralizing Antibody Signatures and Application to Epitope-Targeted Vaccine Design. Cell Host Microbe 25:59–72.e8.

24. Herschhorn A. 2022. Indirect Mechanisms of HIV-1 Evasion from Broadly Neutralizing Antibodies In Vivo. ACS Infect Dis 10.1021/acsinfecdis.2c00573.

25. Cervera H, Ratnapriya S, Chov A, Herschhorn A. 2021. Changes in the V1 Loop of HIV-1 Envelope Glycoproteins Can Allosterically Modulate the Trimer Association Domain and Reduce PGT145 Sensitivity. ACS Infect Dis 7:1558–1568.

26. Juelg B, Stephenson KE, Wagh K, Walker-Sperling VE, Hassell T, DeCamp A, Giorgi EE, Koup RA, Mascola JR, Arduino RC, DeJesus E, Yates NL, Seaman M, Korber B, Barouch D. 2022. Viral Escape During Triple Broadly Neutralizing Antibody Therapy Against HIV-1, p. CROI Abstarct 139. In.

27. Bar KJ, Sneller MC, Harrison LJ, Justement JS, Overton ET, Petrone ME, Salantes DB, Seamon CA, Scheinfeld B, Kwan RW, Learn GH, Proschan MA, Kreider EF, Blazkova J, Bardsley M, Refsland EW, Messer M, Clarridge KE, Tustin NB, Madden PJ, Oden K, O’Dell SJ, Jarocki B, Shiakolas AR, Tressler RL, Doria-Rose NA, Bailer RT, Ledgerwood JE, Capparelli EV, Lynch RM, Graham BS, Moir S, Koup RA, Mascola JR, Hoxie JA, Fauci AS, Tebas P, Chun T-W. 2016. Effect of HIV Antibody VRC01 on Viral Rebound after Treatment Interruption. New England Journal of Medicine 375:2037–2050.

28. Bar-On Y, Gruell H, Schoofs T, Pai JA, Nogueira L, Butler AL, Millard K, Lehmann C, Suárez I, Oliveira TY, Karagounis T, Cohen YZ, Wyen C, Scholten S, Handl L, Belblidia S, Dizon JP, Vehreschild JJ, Witmer-Pack M, Shimeliovich I, Jain K, Fiddike K, Seaton KE, Yates NL, Horowitz J, Gulick RM, Pfeifer N, Tomaras GD, Seaman MS, Fätkenheuer G, Caskey M, Klein F, Nussenzweig MC. 2018. Safety and antiviral activity of combination HIV-1 broadly neutralizing antibodies in viremic individuals. 11. Nature Medicine 24:1701–1707.

29. Sneha Ratnapriya, Karunakar Reddy Pothula, Kim-Marie A. Dam, Durgadevi Parthasarathy, Héctor Cervera Benet, Ruth Parsons, Xiao Huang, Salam Sammour, Katarzyna Janowska, Miranda Harris, Shamim Ahmed, Samuel Sacco, Joseph Sodroski, Michael D. Bridges, Wayne L. Hubbell, Priyamvada Acharya, Alon Herschhorn. 2023. Conformational flexibility of HIV-1 envelope glycoproteins modulates transmitted / founder sensitivity to broadly neutralizing antibodies. bioRxiv 2023.09.13.557082.

30. Wei X, Decker JM, Wang S, Hui H, Kappes JC, Wu X, Salazar-Gonzalez JF, Salazar MG, Kilby JM, Saag MS, Komarova NL, Nowak MA, Hahn BH, Kwong PD, Shaw GM. 2003. Antibody neutralization and escape by HIV-1. Nature 10.1038/nature01470.

31. Mazurov D, Herschhorn A. 2023. Ultrasensitive quantification of HIV-1 cell-to-cell transmission in primary human CD4+ T cells measures viral sensitivity to broadly neutralizing antibodies. bioRxiv 10.1101/2023.09.08.556871.

32. Flemming J, Wiesen L, Herschhorn A. 2018. Conformation-Dependent Interactions Between HIV-1 Envelope Glycoproteins and Broadly Neutralizing Antibodies. AIDS research and human retroviruses 34:794–803.

33. Yen P-J, Herschhorn A, Haim H, Salas I, Gu C, Sodroski J, Gabuzda D. 2014. Loss of a conserved N-linked glycosylation site in the simian immunodeficiency virus envelope glycoprotein V2 region enhances macrophage tropism by increasing CD4-independent cell-to-cell transmission. Journal of virology 88:5014–28.

34. Myers G, Lenroot R. 1992. HIV Glycosylation: What Does It Portend? AIDS Research and Human Retroviruses 8:1459–1460.

35. Seabright GE, Cottrell CA, van Gils MJ, D’addabbo A, Harvey DJ, Behrens AJ, Allen JD, Watanabe Y, Scaringi N, Polveroni TM, Maker A, Vasiljevic S, de Val N, Sanders RW, Ward AB, Crispin M. 2020. Networks of HIV-1 Envelope Glycans Maintain Antibody Epitopes in the Face of Glycan Additions and Deletions. Structure 28:897–909.e6.

36. Behrens A-J, Crispin M. 2017. Structural principles controlling HIV envelope glycosylation. Curr Opin Struct Biol 44:125–133.

37. Mouquet H, L S, Z E, Y L, C E, JF S, A H-S, PN G, DI S, MS S, H S, T F, MC N, PJ B. 2012. Complex-type N-glycan recognition by potent broadly neutralizing HIV antibodies. 47. Proceedings of the National Academy of Sciences of the United States of America 109.

38. Pancera M, Yang Y, Louder MK, Gorman J, Lu G, McLellan JS, Stuckey J, Zhu J, Burton DR, Koff WC, Mascola JR, Kwong PD. 2013. N332-Directed broadly neutralizing antibodies use diverse modes of HIV-1 recognition: inferences from heavy-light chain complementation of function. PloS one 8:e55701.

39. Shingai M, Donau OK, Schmidt SD, Gautam R, Plishka RJ, Buckler-White A, Sadjadpour R, Lee WR, LaBranche CC, Montefiori DC, Mascola JR, Nishimura Y, Martin MA. 2012. Most rhesus macaques infected with the CCR5-tropic SHIVAD8 generate cross-reactive antibodies that neutralize multiple HIV-1 strains. Proceedings of the National Academy of Sciences 109:19769–19774.

40. Francica JR, Sheng Z, Zhang Z, Nishimura Y, Shingai M, Ramesh A, Keele BF, Schmidt SD, Flynn BJ, Darko S, Lynch RM, Yamamoto T, Matus-Nicodemos R, Wolinsky D, Nason M, Valiante NM, Malyala P, De Gregorio E, Barnett SW, Singh M, O’Hagan DT, Koup RA, Mascola JR, Martin MA, Kepler TB, Douek DC, Shapiro L, Seder RA, Barnabas B, Blakesley R, Bouffard G, Brooks S, Coleman H, Dekhtyar M, Gregory M, Guan X, Gupta J, Han J, Ho SL, Legaspi R, Maduro Q, Masiello C, Maskeri B, McDowell J, Montemayor C, Mullikin J, Park M, Riebow N, Schandler K, Schmidt B, Sison C, Stantripop M, Thomas J, Thomas P, Vemulapalli M, Young A. 2015. Analysis of immunoglobulin transcripts and hypermutation following SHIVAD8 infection and protein-plus-adjuvant immunization. Nature Communications 6.

41. Walker LM, Huber M, Doores KJ, Falkowska E, Pejchal R, Julien JP, Wang SK, Ramos A, Chan-Hui PY, Moyle M, Mitcham JL, Hammond PW, Olsen OA, Phung P, Fling S, Wong CH, Phogat S, Wrin T, Simek MD, Koff WC, Wilson IA, Burton DR, Poignard P. 2011. Broad neutralization coverage of HIV by multiple highly potent antibodies. 7365. Nature 477:466–470.

42. Kielian M, Rey FA. 2006. Virus membrane-fusion proteins: more than one way to make a hairpin. 1. Nat Rev Microbiol 4:67–76.

43. Rey FA, Lok S-M. 2018. Common Features of Enveloped Viruses and Implications for Immunogen Design for Next-Generation Vaccines. Cell 172:1319–1334.

44. Herschhorn A, Ma X, Gu C, Ventura JD, Castillo-Menendez L, Melillo B, Terry DS, Smith AB, Blanchard SC, Munro JB, Mothes W, Finzi A, Sodroski J. 2016. Release of gp120 Restraints Leads to an Entry-Competent Intermediate State of the HIV-1 Envelope Glycoproteins. mBio 7:e01598–16.

45. Herschhorn A, Gu C, Moraca F, Ma X, Farrell M, Smith AB, Pancera M, Kwong PD, Schön A, Freire E, Abrams C, Blanchard SC, Mothes W, Sodroski JG. 2017. The β20-β21 of gp120 is a regulatory switch for HIV-1 Env conformational transitions. Nature Communications 8:1049.

46. Wang K, Zhang S, Go EP, Ding H, Wang WL, Nguyen HT, Kappes JC, Desaire H, Sodroski J, Mao Y. 2023. Asymmetric conformations of cleaved HIV-1 envelope glycoprotein trimers in styrene-maleic acid lipid nanoparticles. Commun Biol 6:535.

47. Harris M, Ratnapriya S, Chov A, Cervera H, Block A, Gu C, Talledge N, Mansky LM, Sodroski J, Herschhorn A. 2020. Slow Receptor Binding of the Noncytopathic HIV-2UC1 Envs Is Balanced by Long-Lived Activation State and Efficient Fusion Activity. Cell Rep 31:107749.

48. Ratnapriya S, Chov A, Herschhorn A. 2020. A Protocol for Studying HIV-1 Envelope Glycoprotein Function. STAR Protoc 1:100133.

49. Herschhorn A, Marasco W a, Hizi A. 2010. Antibodies and lentiviruses that specifically recognize a T cell epitope derived from HIV-1 Nef protein and presented by HLA-C. Journal of immunology 185:7623–32.

50. Herschhorn A, Hizi A. 2008. Virtual screening, identification, and biochemical characterization of novel inhibitors of the reverse transcriptase of human immunodeficiency virus type-1. J Med Chem 51:5702–5713.

51. Weitman M, Lerman K, Nudelman A, Major DT, Hizi A, Herschhorn A. 2011. Structure-activity relationship studies of 1-(4-chloro-2,5-dimethoxyphenyl) -3-(3-propoxypropyl)thiourea, a non-nucleoside reverse transcriptase inhibitor of human immunodeficiency virus type-1. European Journal of Medicinal Chemistry 46:447–467.

52. Ratnapriya S, Braun AR, Cervera Benet H, Carlson D, Ding S, Paulson CN, Mishra N, Sachs JN, Aldrich CC, Finzi A, Herschhorn A. 2022. Broad Tricyclic Ring Inhibitors Block SARS-CoV-2 Spike Function Required for Viral Entry. ACS Infect Dis 8:2045–2058.

53. Herschhorn A, Oz-Gleenberg I, Hizi A. 2008. Mechanism of inhibition of HIV-1 reverse transcriptase by the novel broad-range DNA polymerase inhibitor N-{2-[4-(aminosulfonyl)phenyl]ethyl}-2-(2-thienyl)acetamide. Biochemistry 47:490–502.

54. Ratnapriya S, Harris M, Chov A, Herbert ZT, Vrbanac V, Deruaz M, Achuthan V, Engelman AN, Sodroski J, Herschhorn A. 2021. Intra- and extra-cellular environments contribute to the fate of HIV-1 infection. Cell Rep 36:109622.

55. Herschhorn A, Finzi A, Jones DM, Courter JR, Sugawara A, Smith AB, Sodroski JG. 2011. An inducible Cell-Cell fusion system with integrated ability to measure the efficiency and specificity of HIV-1 entry inhibitors. PLoS ONE 6.

56. Meng EC, Goddard TD, Pettersen EF, Couch GS, Pearson ZJ, Morris JH, Ferrin TE. 2023. UCSF ChimeraX: Tools for structure building and analysis. Protein Sci 32:e4792.

